# Latitude-Driven Functional Trait Variations in *Zizania latifolia*: Insights into Climate Adaptation

**DOI:** 10.1101/2023.10.29.564516

**Authors:** Hongsheng Jiang, Xiangrong Fan, Godfrey Kinyori Wagutu, Wei Li, Yuanyuan Chen

## Abstract

The global warming could have significant impact on plant adaptation to local environments. Widespread species provides a useful model to examine the population dynamic under climate change. However, it is still unclear how widespread aquatic species response to rapidly changing environments. Wild-rice *Zizania latifolia* is an emergent macrophyte widely distributed across East Asia. Here, 28 *Z. latifolia* natural populations covering above 30 latitudes were transplanted in a common garden. The growth, photosynthesis and reproduction traits were quantified and compared among populations, and pairwise relationships among geographic and genetic variables were estimated. Results showed that, in the common garden, the high-latitude populations grew in small size indicating weaker competitiveness in warmer temperatures; the low-latitude populations had no sexual reproduction, suggesting that low-latitude individuals stood little chance to migrate successfully to higher latitude. Significant positive correlations among the distances of genetic, geographic and morphological traits for populations indicated that both isolation by distance and isolation by environment models affected the genetic pattern of populations, and phenotypic traits of *Z. latifolia* populations might be genetically determined. All our data suggest that *Z. latifolia* would be inadaptable to the global warming, indicating that this species would be at least at risk of local extinction in the warmer future.

## INTRODUCTION

The successful establishment of plant populations is most commonly attributed to the well-adapted individual genotypes, which have been suitable to the local environmental conditions, and shaped over a long evolutionary history (Joshi et al., 2001). However, the accelerated global environmental changes, represented by global warming, have greatly impacted the species adaption potential (Cox et al., 2000). Global warming will reduce the fitness of many wild plants in their existing natural habitats. These species can migrate to cooler habitats, like the areas with high altitudes or latitudes, by dispersing spores, seeds and/or propagules (Kelly and Goulden, 2008; Lippmann et al., 2019). The survival of migratory species depends on how effectively it can compete with the native species of the novel environments. Alternatively, plant species can adapt to climate change through natural selection, a slow process dependent on random mutations. However, it was predicted that natural selections in many species can’t keep pace with the global warming and is insufficient for rapid adaptation (Wilczek et al., 2014). For plants that remain in their original habitats, global warming causes them to undergo morphological and phenological changes during their life histories. A general response of plants to temperature rise is to grow taller and slender with leaves narrower and thinner, which will minimize body temperature of plants (Lee et al., 2020). Notably, these morphogenic changes cause plants to bend easily in the wind and rain, and result in a loss of biomass (Mooney et al., 2010). The most studied phenological changes are the changes in flowering time, and as a result, pollinators are unable to pollinate efficiently (Borghi et al., 2019; Lippmann et al., 2019). Besides the direct impact on plant growth and reproduction within eco- and agricultural systems, the changing environments have increased the vulnerability of plants to attack by different pathogens and pests(Cohen and Leach, 2020; Desaint et al., 2021; Hamann et al., 2021). Therefore, under global warming, species that either move to new places or remain in their native habitats are predicted to become extinct. It is notable that most previous studies about plant response to temperature increase are from the model plant *Arabidopsis thaliana* or the cultivated plants, especially major staple crops (Lippmann et al., 2019). Although some temperature responses are conserved across species, most responses vary greatly within and among different plant species (Kim et al., 1996; Commuri and Jones, 2001). Furthermore, phenological events, like flowering and autumnal growth cessation, are controlled by both temperature and photoperiod (Korner and Basler, 2010). Therefore, when plants migrate to higher latitudes due to global warming, they will face new challenges from photoperiodic changes. Global temperature has been expected to increase 1.1-6.4 °C at the end of this century (Solomon, 2007). As a response, the plant extinction rate is predicated to be 1 000 to 10 000 times higher than that of the background (Pimm and Joppa, 2015; Levin, 2019). Therefore, the urgent situation calls for a better understanding of plant responses to global warming, especially for economically and/or ecologically important species.

Functional traits are characterized by the response of morphology, physiology and phenology to the environment, and to the interaction among species (Díaz and Cabido, 2001; Violle et al., 2007). Compared to individual analysis, functional traits make it possible to analyze and compare plant functioning at ecosystem, landscape, or regional scale, because the entity plant individuals are replaced by a set of observable traits (Soudzilovskaia et al., 2013). Therefore, functional traits have been used to describe life systems in different levels for various aims, such as in levels of population, community, ecosystem and global vegetation (Heilmeier, 2019). Widespread plant species are composed of various genotypes or possess high phenotypic plasticity, and their functional traits can respond differently to climate change because of the adaption to local climate conditions (Giudicelli et al., 2019; Ren et al., 2020). Such plants thus provide a useful model to understand the variation of functional traits under climate changes, and evaluate the species destiny. A common garden experiment can effectively assess whether the plants acclimate or adapt to the climate changes when the populations from different climatic regions are transplanted in a common environment (Xiankui and Chuankuan, 2018). Due to the possible adaptation to the various environmental conditions, the widespread species possess high morphological and/or genetic variations. A common garden experiment is thus an ideal approach to evaluate if the intraspecific trait variations are genetically based or phenotypic plasticity (Ren et al., 2020). Because of these advantages, many common garden experiments have been conducted in researches about plant responses to climate change associated with altitude and latitude shifting (Vitasse et al., 2009; Buckley et al., 2019; Ren et al., 2020).

Wild rice *Zizania* L., a widespread aquatic angiosperm genus from the tribe *Oryzeae* (Poaceae), comprises one species distributed in East Asia (*Z. latifolia*), and three species from North America (*Z. aquatica*, *Z. palustris* and *Z. texana*) (Terrell et al., 1997; Guo et al., 2007; Porter, 2019). Among these four species, *Z. palustris* and *Z. latifolia* are of important agricultural values as aquatic crops (Oelke, 1993; Guo et al., 2007; Porter, 2019). *Z. latifolia*, known as Manchurian wild rice, is a perennial, monoecious, emergent aquatic plant. It occurs in littoral of freshwater streams and marshes and shallow water of lake margins and swamps, and is widely distributed across China. *Z. latifolia* is wind-pollinated and reproduces asexually by rhizomes or sexually by seeds. It served as an important grain in ancient China and has been a popular aquatic vegetable in the last 1500 years in form of swollen young shoots after being infected by smut fungus (Guo et al., 2007). The eminent breeding traits of *Z. latifolia* have also been used to improve rice varieties due to its close phylogenetic relationship to rice (*Oryza sativa*)(Fan et al., 2016; Zhao et al., 2018). Besides the economic values, wild populations of *Z. latifolia* have important ecological values, including strengthening levees and purifying water, properties associated with their powerful clonal reproduction and strong nutrient absorption capacity (Guo et al., 2007). Thus, obtaining knowledge about the functional trait variations of *Z. latifolia* under climate changes could not only help us to better understand the widespread species population dynamics, but also benefit the formulation of management strategy for the important agricultural resource.

In the present study, the East Asia widespread species, *Z. latifolia* from gradient latitudes (N 20-50°) was chosen as a model plant to estimate growth and reproduction traits changes under global climate change in a common garden (N 30°). With these investigations, we attempt to address: 1) how functional traits of *Z. latifolia* from different latitudes change in the common garden; 2) how the temperature affects the functional traits; and 3) whether *Z. latifolia* could adapt to a warming future.

## MATERIALS AND METHODS

### Sampling and Climate Data Collection

In the Autumn of 2015, 28 *Z. latifolia* populations were collected from five regions with gradient latitude (N 20°21′ - 50°54′), including Heilongjiang River basin (HLJ), Liaohe River basin (LHR), Huanghe River basin (HHR), Yangtze River basin (YZR) and Zhujiang River basin (ZJR), covering more than 30 latitudes across the north to south of China (Table S1 and Figure 1). For each river basin, four to seven separated populations were randomly collected. In each population, more than 20 well-grown individuals, approximately 1.8 m in height above ground or water surface, were harvested with intervals at least 10 m to minimize the opportunity of sampling the same clone. In all, 580 individuals were collected from the 28 populations. Healthy stems (about 1 m in length) and young leaves were collected for each individual. The leaves were immediately place in silica-containing plastic bags, and the stems were planted in Wuhan Botanical Garden (N 30.544, E 114.420). The individuals were genotyped by 25 SSR (simple sequence repeat, or microsatellite) markers (Table S2). For each population, the identified clones deleted from further experiments.

**Figure 1.**
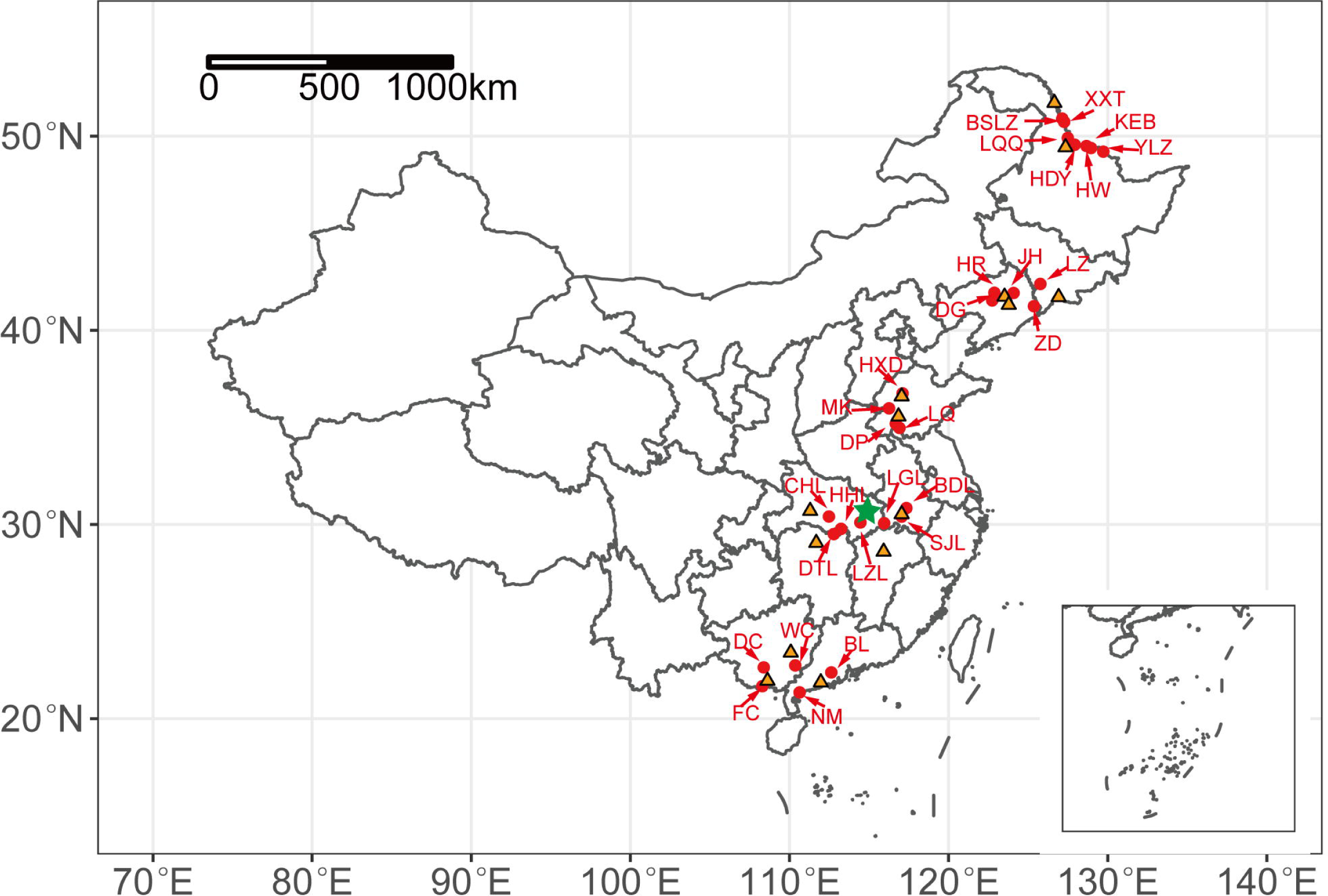
Sampling sites of *Zizania latifolia* populations and location of meteorological stations. The red circle points represent *Z. latifolia* populations, the orange triangle points represent meteorological stations, the green star indicates the location of the common garden.

Climate data was extracted from National Oceanic and Atmospheric Administration (NOAA). The daily temperature and precipitation values across 60 years (1954 to 2014) were obtained from 14 meteorological stations nearby our sampling sites (Fig 1 and Table S3; https://gis.ncdc.noaa.gov/maps/ncei/summaries/daily). Two or three populations could share the same meteorological data when the geographic distance between or among the sites are less than 100 km.

### Common Garden Experiments

To investigate phenotypic variation in populations within and between latitude regions and its correlation with habitats, a common garden was established using the 28 populations from five river basins across China. In April of 2017, except for population XXT which had only 10 surviving individuals, 14 randomly selected genets of each population were grown to the tillering stage in Wuhan Botanical Garden. Then, we chose three ramets of each plant, and all ramets were of similar size, with three leaves and about 30 cm in height. In all, the 1164 ramets were planted into pots (50 cm in diameter, 32 cm in height) and transplanted into a common garden located in Wuhan suburban (N 30.730; E 114.165). The *Z. latifolia* plantlet pots were arranged in rows with 1.0 m between the rows and 0.5 m between pots at the experimental area (40 × 40 m^2^). Daily management of the common garden was carried out by local farmers.

### Growth and Reproduction Traits

The heading date of each population was recorded for each population from August to December. In August, the highest tiller of each plant was chosen to measure plant height, leaf length (of the longest leaf), leaf width (of the longest leaf), internode number, internode length, and stem diameter. In December, the number of tillers and spikes and the height of spikes were recorded; the number of seeds for each spike was measured by counting the ends of the spike branches because of the seed shattering habit of *Z. latifolia*. We also calculated the frequency of plant spike. It was counted as ‘1’ if the individual bloomed, or it was counted as ‘0’ for the individual without bloom. Then, the spike relative frequency was calculated of each population (the number of bloomed individuals divided by the total individual number of population).

After harvesting, the aboveground part of each plant was dried in oven at 80 °C for three days, and weighed for biomass estimation.

### Photosynthesis and Respiration Traits

Ten individuals in each population of LQQ, HR, MK, DT and BL were randomly chosen to represent the five latitude regions (HLJ, LHR, HHR, YZR and ZJR) for photosynthesis and respiration traits measurement, including chlorophyll fluorescence, photosynthetic rate and respiration rate. Chlorophyll fluorescence parameters was determined by a “Monitoring-PAM Multi-Channel Chlorophyll Fluorometer” (Walz, Effeltrich, Germany). Briefly, after 15 min dark adaptation, the maximum quantum yield of PSII (Fv/Fm) of *Z. latifolia* leaf was calculated from the minimum fluorescence (Fo) under a measuring light (2 µmol photons m^-2^ s^-1^) and the maximum fluorescence (Fm) under a saturating plus light (3500 µmol photons m^-2^ s^-1^) (Equation 1).

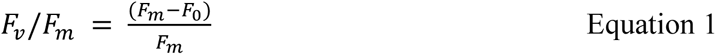

The rapid light curve (RLC) of *Z. latifolia* leaf describes the relationship between relative electron transport rate (rETR) and photosynthetic active radiation (PAR) under increasing illumination using the Equation 2, where rETR_max_ is the maximum potential rETR, α is the RLC initial slope.

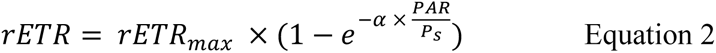

The light-saturation coefficient (Ek) was calculated from RLC using the Equation 3.

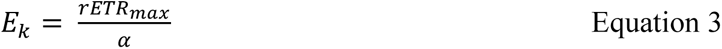

Photosynthesis rate was recorded with a “Li-cor 6400” (Li-Cor Inc., Lincoln, NE, USA) under the ambient CO_2_ concentration (approximately 400 ppm), 1000 µmol photons m^-2^ s^-1^ light intensity and 30 ℃ for 5 min. After photosynthesis rate measurement, the light was turned off, and the respiration rate was recorded.

### Geographic Distance, Genetic Distance and Growth Traits Distance

Based on geographical coordinates, we calculated the geographic distance among populations using the “distVincentyEllipsoid” function in the “geosphere” package (Hijmans) with R program 4.0(Team, 2013). Genetic distance (Nei, 1972) and genetic differentiation among populations (*F*_ST_)(Weir and Cockerham, 1984) was estimated using the software GenAlEx 6.5(Peakall and Smouse, 2006). Trait differentiation (*Q*_ST_) was estimated by the variance among populations (*V*_among_) and the average variance within population (*V*_within_) as following equation (Leinonen et al., 2013):

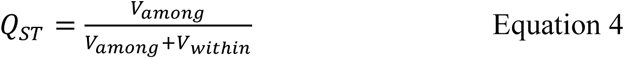

The growth traits data were scaled using the “scale” function in R program. Then, the growth traits distance matrix was determined by Euclidean distance. Due to absence of blooming for individuals in ZJR region in the common garden, the relationships between reproduction traits and geographic or genetic distance were not estimated.

### Statistical Analysis

Using R program 4.0(Team, 2013), a linear model was used to fit the trends of yearly average temperature and annual precipitation in locations of the *Z. latifolia* populations distribution from the year 1954 to 2014. Non-parameteric Kruskal-Wallis multiple comparison with *P*- values adjusted using the “Holm” method (*P* < 0.05) was performed to determine the differences of growth traits and reproduction traits among sampling regions (HLJ, LHR, HHR, YZR and ZJR), because the data do not follow a normal distribution. Depending on the R square of regression model, one of the three models: 1) a linear model without interaction, 2) a linear model with interaction, or 3) a binomial regression model, was chosen to determine relationship between growth and reproduction traits of *Z. latifolia* in the common garden and their original habitat yearly average temperature and annual precipitation. A linear mixed model was chosen to determine differences of the rapid light curves (RLC) among sampling regions by using R program “nlme” package, where sampling region and photosynthetic active radiation (PAR) were set as two fixed factors and the plant individual was set as a random factor. One-way ANOVA followed by Tukey test were performed to determine differences of RLC initial slope (α), light-saturation coefficient (Ek), photosynthesis rate, and respiration rate among sampling regions by in R program. Cluster analysis was conducted within growth traits using “mclust” package (Scrucca et al., 2016) and “pheatmap” using the “Euclidean distance” after applying “scale” to the data. The reproduction traits were excluded from the cluster analysis because there were no individuals that bloomed in the ZJR region. A linear model was chosen to determine the relationships among geographic distance, genetic distance and growth traits distance, with square or square root transformation performed if necessary.

## RESULTS

### Sampling Site Climate Change

The logs from the 14 selected meteorological stations showed that the yearly average temperature went up steadily in the past 60 years (Figure 2). The most rapid temperature increment was found in the northeast sampling region, HJL, which was up to 5.4 ℃ per century. For the other four regions, LHR, HHR, YZR, and ZJR, the yearly average temperature increased in the range of 1.1 to 2.5 ℃ per century. In contrast to the temperature, no significant trend was found in the annual precipitation across all sampling regions (Figure S1).

**Figure 2.**
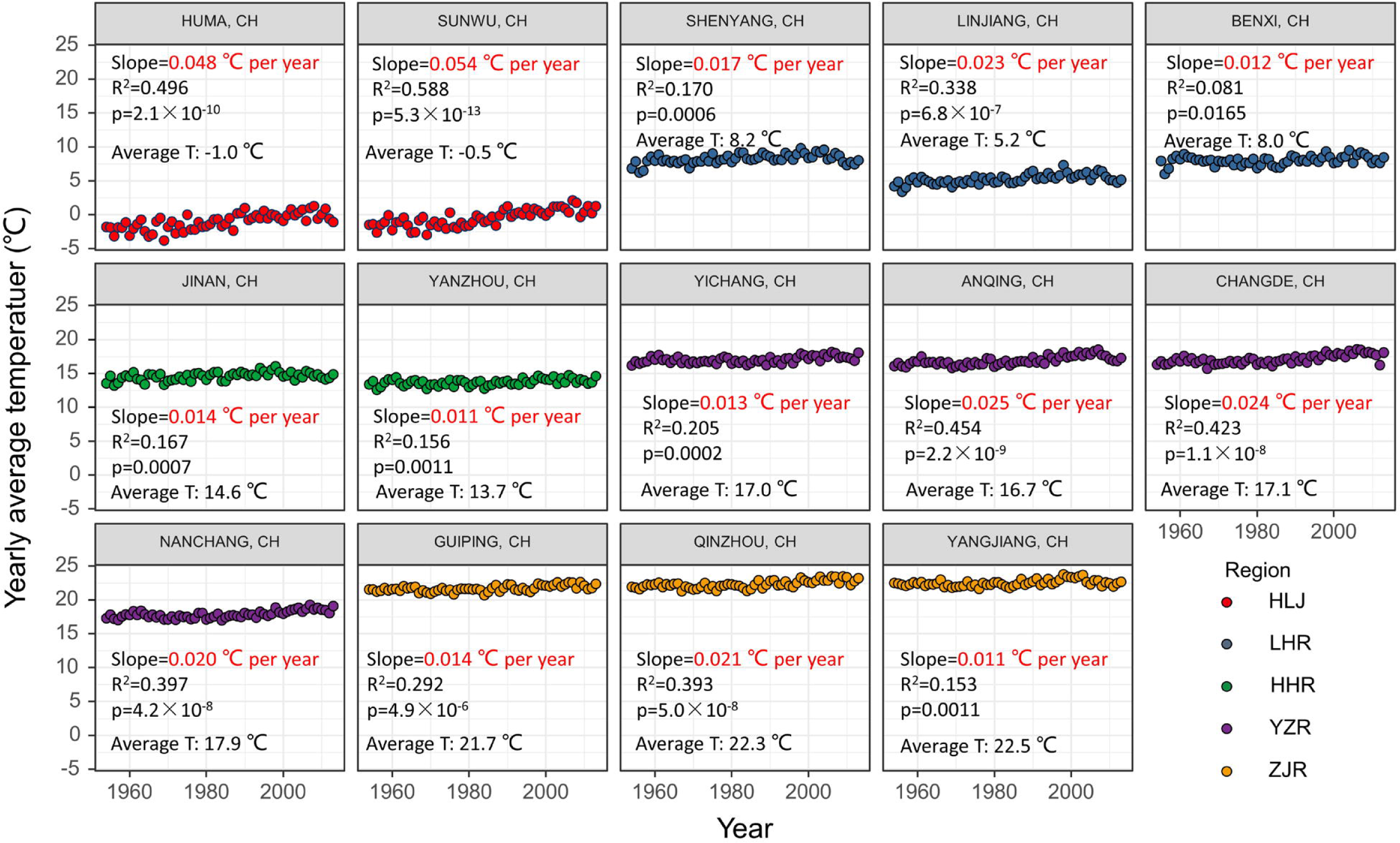
Changes of yearly average temperature at sampling sites in the past 60 years (1954-2014).

### Growth Traits

In the common garden, most growth traits were significantly different among the sampling regions. The most and the second most tiller number were observed in the regions of LHR and of YZR (Figure 3a). Both temperature and precipitation of original habitat significantly affect plant tiller number, which could explain 36% variation of tiller number in the common garden. Either too high or too low temperature decreased tiller number (Figure 3b and Table S4). There were significant differences in plant height (Figure 3c), leaf length (Figure 3e), and leaf width (Figure 3g) among regions. Compared with those from the north, the plants that originated from the south grew longer and wider leaves and taller plant heights, which could have been due to the higher temperature and precipitation in the original habitat (Figure 3d, f, and h). The two climate factors could explain more than 85% variation of plant height, leaf length and leaf width of *Z. latifolia* (Table S4). However, the populations that originated from the south China (ZJR region) had significantly less internode number and significantly shorter internode length (Figure 3i and k). The internode number and internode length were significantly correlated to temperature and precipitation of original habitats, despite less than 20% variation that could be explained (Figure 3j and l, Table S4). Original habitat temperature and precipitation explained high level of variations in the plant dry weight (95%) and stem diameter (76%) (Figure 3n and p; Table S4), which showed an increasing pattern from the south to north in the common garden (Figure 3m and o). Based on all growth traits, the cluster analysis showed that all *Z. latifolia* populations clustered into two groups (Figure 3q): (1) group I contained the populations from the three regions, HHR, YZR and ZJR; (2) group II contained the populations from the regions HLJ and LHR.

**Figure 3.**
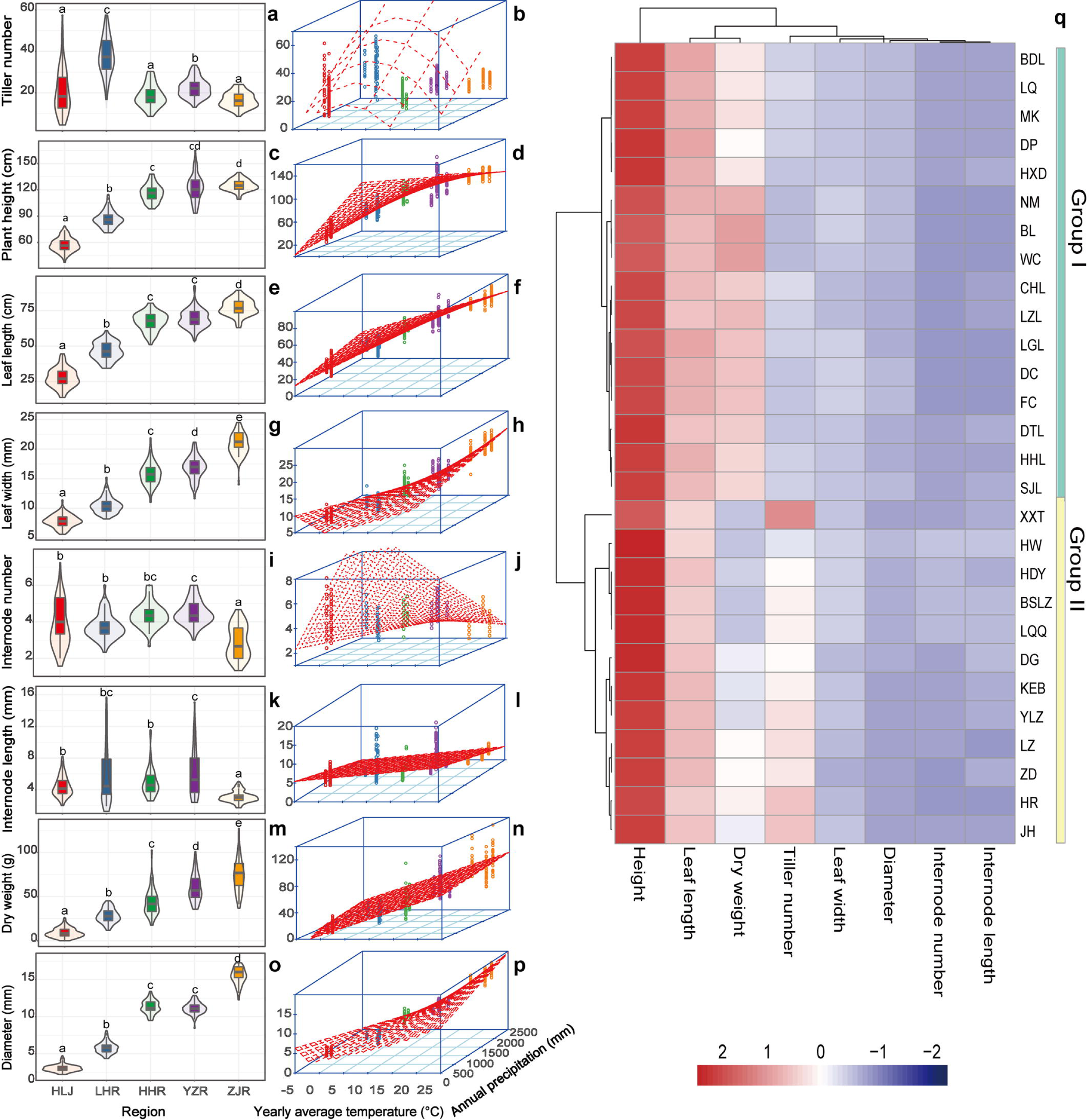
Differences of growth traits of *Zizania latifolia* among sampling regions and their relationships between climate factors, and the cluster analysis of growth traits within populations. (a) difference of tiller number among sampling regions; (b) relationship between tiller number and climate factors; (c) difference of plant height among sampling regions; (d) relationship between plant height and climate factors; (e) difference of leaf length among sampling regions; (f) relationship between leaf length and climate factors; (g) difference of leaf width among sampling regions; (h) relationship between leaf width and climate factors; (i) difference of internode number among sampling regions; (j) relationship between internode number and climate factors; (k) difference of internode length among sampling regions; (l) relationship between internode length and climate factors; (m) difference of dry weight among sampling regions; (n) relationship between dry weight and climate factors; (o) difference of stem diameter among sampling regions; (p) relationship between stem diameter and climate factors; and (q) cluster analysis of growth traits in the 28 populations.

### Photosynthesis Traits

Rapid light curve (RLC) showed no significant difference of photosynthesis capacity among sampling regions (*F* = 0.976, *P* = 0.4306, Figure 4a and b). The three parameters α (Figure 4c), E_k_ (Figure 4d), and rETR_max_ (Figure 4e) also showed no significant difference among sampling regions. Similarly, the maximum quantum yield of photosystem II (Fv/Fm) and photosynthesis rate did not significantly change with the sampling regions (Figure 4f and g). In contrast, significantly higher respiration rate (Figure 4h) was observed in the north sampling regions (HLJ and LHR region) than that in the south sampling regions (HHR, YZR and ZJR region).

**Figure 4.**
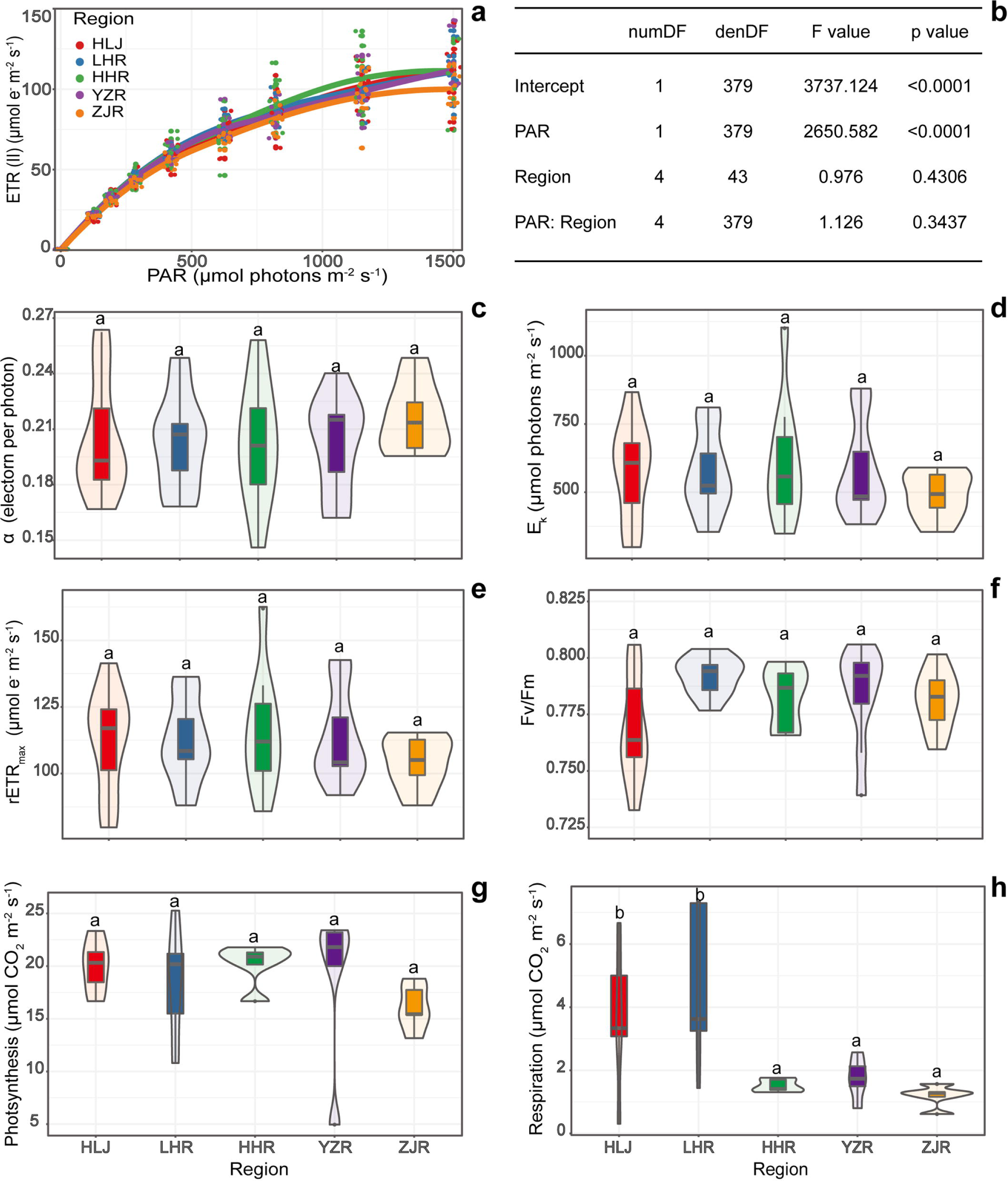
Photosynthesis traits of *Zizania latifolia* among sampling regions. (a) rapid light curve (RLC) among sampling regions, (b) statistical analysis result of RLC; following parameters of RLC: α (c), Ek (d), and rETRmax (e); (f) maximum quantum yield of photosystem II (Fv/Fm); (g) photosynthesis rate and (h) respiration rate.

### Reproduction Traits

More than 70% individuals bloomed in the north originated populations (HLJ, LHR and HHR). However, only approximately 10% individuals bloomed in the YZR region populations, and no individual bloomed in populations that originated from the south-most region (ZJR; Figure 5a). The relative frequency of spike was significantly determined by original habitat temperature which explained approximately 60% variation of the spike frequency among various regions (Figure 5b and Table S4). Compared with those from the south regions, *Z. latifolia* originating in north regions (HLJ, LHR and HHR) showed higher blooming frequency and more spikes per individual (Figure 5c). Too high or too low temperature reduced the number of spikes per individual and higher precipitation promoted spike development (Figure 5d). The *Z. latifolia* individuals from the south had a higher spike and more grain number per spike than those of the plants from north, except for the individuals from the south-most region (ZJR), which didn’t bloom in the common garden (Figure 5e and g). Higher temperature in original habitat could increase spike length and it could explain 76% variation of spike length among regions, however precipitation had no significant effects on the spike length (Table S4). Original habitat temperature and precipitation significantly affected the grain number per spike, which determined 63% variation of grain number per spike (Figure 5 h and Table S4).

**Figure 5.**
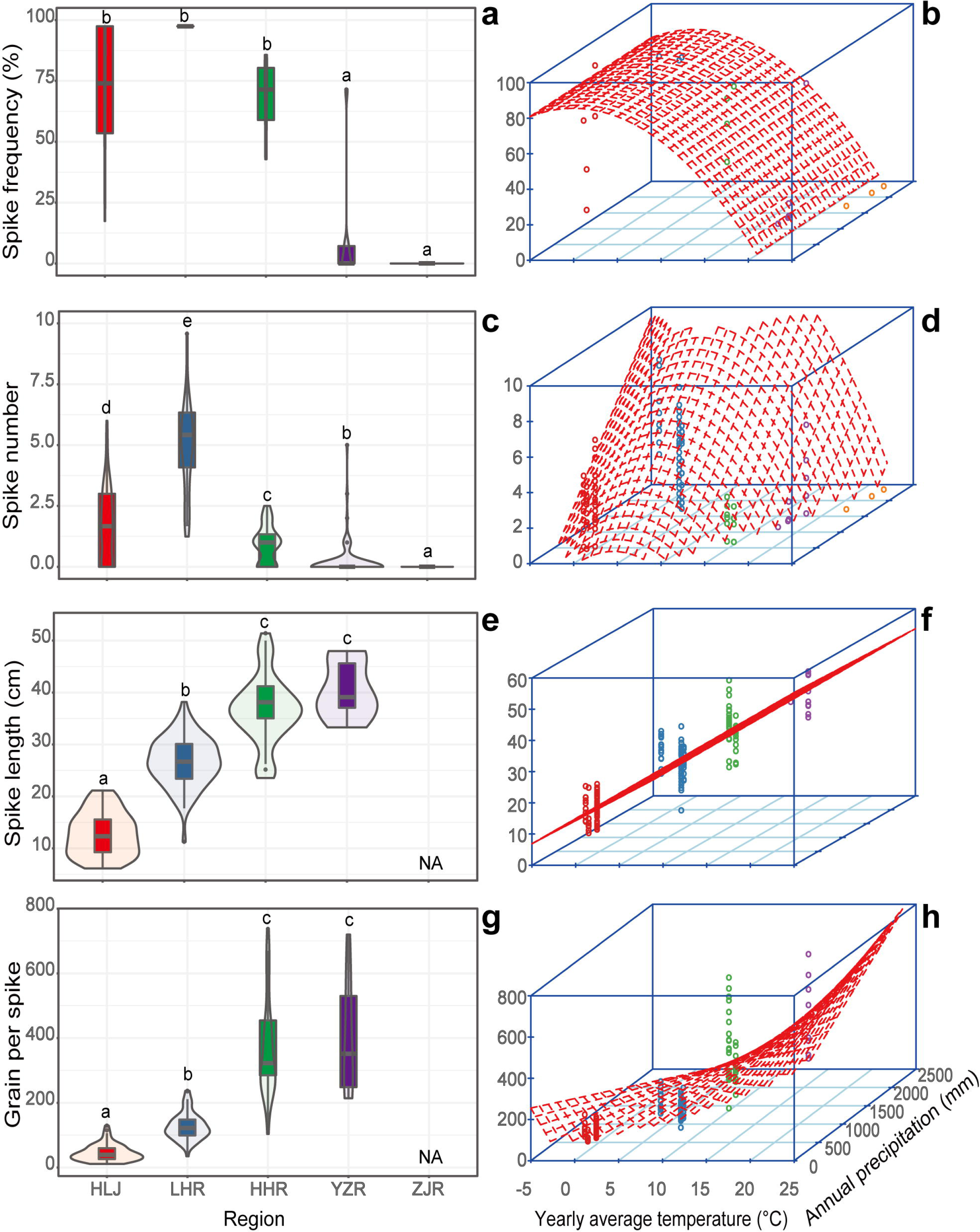
Differences of reproduction traits of *Zizania latifolia* among sampling regions and their relationships between climate factors, and the cluster analysis of growth traits within populations. (a) difference of relative spike frequency among sampling regions; (b) relationship between relative spike frequency and climate factors; (c) difference of spike number per individual among sampling regions; (d) relationship between spike number per individual and climate factors; (e) difference of spike length among sampling regions; (f) relationship between spike length and climate factors; (g) difference of grain per spike among sampling regions; (h) relationship between grain per spike and climate factors.

### Relationships among Geographic, Genetic and Growth Traits Distance

The global genetic differentiation across all populations was suggested by *F*_ST_ = 0.585 (*P* = 0.001) and all the trait differentiation (*Q*_ST_ = 0.931-0.998; Table S5) was higher than *F*_ST_. Genetic distance (Nei, 1972) increased with the increasing geographic distance (Figure 6a) and growth trait distance increased with the increasing genetic distance (Figure 6b). A high correlation (*r*^2^ = 0.716) was observed between growth distance and geographic distance (Figure 6c).

**Figure 6.**
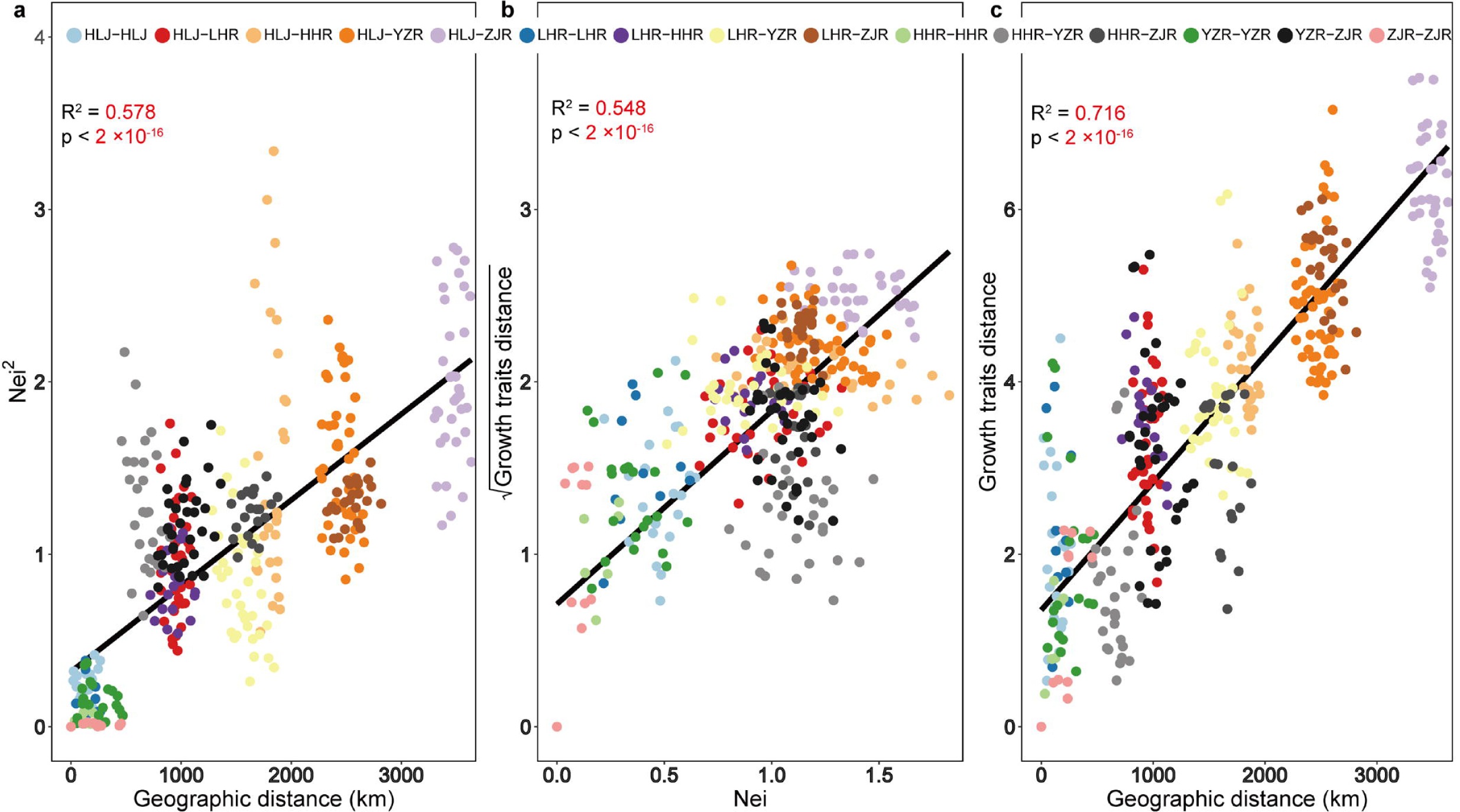
Relationships among geographic distance, genetic distance (Nei) and growth traits distance of *Zizania latifolia* among sampling sites. (a) relationship between genetic distance and geographic distance; (b) relationship between growth distance and genetic distance; (c) relationship between growth traits distance and geographic distance.

## DISCUSSION

Climate-mediated natural selection is one of processes that drive local adaptation and speciation (Lv et al., 2014). Previous studies have reported that the annual average temperature would continue to increase, which was the case in all the current sampling sites, especially in the northeast China (Figure 2). According to the trend, we predict that the temperature in the northeast China will increase by 5℃ per century, which is comparable to the prediction from Li et al. (2020) and IPCC (Solomon, 2007). Therefore, in future decades, the temperature in HLJ region will reach 5 ℃, which is equal to the current temperature (∼7 ℃) at LHR region. Furthermore, in future centuries, it could even increase to 15∼20℃, equivalent to the current temperature in south China.

### Growth, Photosynthesis and Respiration Traits

Although both the yearly average temperature and annual precipitation significantly affected the growth traits of *Z. latifolia*, we will focus on the temperature effects in the following discussion considering no significant change in the annual precipitation within each region in the past 60 years (Figure S1).

In the common garden, compared with the populations from the three low-latitude regions (HHR, YZR and ZJR), the individuals of populations from the two high-latitude regions (HLJ and LHR) showed a small and compact plant type, with shorter height and smaller leaves (Figure 3). Additionally, with the annual temperature increasing at 1 ℃, the plant height, leaf length and width would decrease by 4.65 cm, 2.81 cm and 0.39 mm (Table S4), respectively. Previous studies showed that the individuals with taller height and larger leaves had more powerful competitiveness because of their advantages in harvesting light, the source of primary productivity(Falster and Westoby, 2003; Craine et al., 2013). Therefore, the compact plant type of the populations from high latitude might suggest the downregulated competitiveness when temperature increases.

Plant architecture, such as the number and length of internode, represents plant mechanical supporting and resource allocation (Niklas, 1994; Messier et al., 2017). In the present study, compared with YZR populations which shared the same climate conditions in the common garden and in wild, the higher- and lower-latitude individuals had shorter internode length and fewer internode numbers, indicating the decrease of resource allocation in internodes. Furthermore, the low-latitude individuals showed larger stem diameter and more biomass. The results suggested high competitiveness in low latitude populations because large-diameter plants contained more biomass and carbon storage which played disproportionate roles to ecosystem function (Lutz et al., 2013).

The warming temperature could increase (Wan et al., 2002; Niu and Wan, 2008) or decrease (Hobbie et al., 1999) plant productivity which was often linked to biomass accumulation. In this study, warming induced *Z. latifolia* biomass decrease by showing the lighter dry weight of high-latitude populations. Photosynthesis, as the process of synthesizing organic matter, is generally considered as the most important factor for biomass accumulation (Raines, 2011; Chen et al., 2018). However, all *Z. latifolia* populations showed similar levels of photosynthetic parameters, including rapid light curve (RLC), maximum electron transport (rETRmax), light-saturation coefficient (Ek), maximum quantum yield (Fv/Fm), and photosynthesis rate. Alternatively, respiration, the process of consuming organic matter, also impacts on biomass accumulation. In this study, higher respiration rate was found in the high-latitude regions compared with the low-latitude regions. Therefore, instead of photosynthesis, the respiration dominated the biomass accumulation in the common garden. Respiration often reaches to the peak at a higher temperature than that for photosynthesis (Atkin, 2003). This could result in a decrease of biomass accumulation due to higher organic matter degradation than synthesis under a warmer condition (Atkin et al., 2007; Smith and Dukes, 2013).

The growth trait changes of *Z. latifolia* driven by temperature were also evident from the cluster analysis. The common garden was located in YZR with a yearly average temperature of ∼17 ℃. In this study, all populations clustered into two groups. For the three regions in group I, the temperature difference between YZR and HHR/ZJR was less than 5 ℃; while for the group II, there was a large temperature difference between YZR and LHR (10 ℃)/ HLJ (18 ℃), and a small temperature difference between LHR and HLJ (8 ℃).

### Reproduction and Photoperiod

Effective sexual reproduction is the foundation for long-term maintenance and regeneration of plant populations, although asexual reproduction could be beneficial for plants to rapidly occupy specific habitats in stable environments (Aguilar et al., 2006; Lively and Morran, 2014). In the present study, the higher-latitude populations produced more spikes than those from the lower latitude regions, indicating that the sexual reproduction increased when the populations were moved to lower latitude. Despite the more bloom opportunities and more number of spikes in higher latitude populations, their shorter spike length and less grain per spike suggested that the less biomass accumulation couldn’t supply enough energy for sexual reproductivity, even when the plant attempted to allocate more resource to sexual reproduction. Although the YZR populations bloomed less, they grew the longer spike and more grains per spike, indicating the effective sexual reproduction in these populations.

Although the effects of photoperiod sometimes could be obscured by temperature and precipitation (Ren et al., 2020), most previous studies showed the importance of photoperiod in plant phenology (refer to the review of Pau et al., 2011). In this study, the photoperiod should have been significantly altered when higher- or lower-latitude individuals were moved to the common garden. There were two main ecotypes in cultivated varieties of *Z. latifolia*: one was a strictly short-day plant harvested in fall; the other one was insensitive to light and harvested in fall and summer (Zhao et al., 2019). Given that the swollen shoots of wild *Z. latifolia* could only be harvested in fall when infected by smut fungus, we inferred that wild *Z. latifolia* was a short-day plant. The short-day plants *Cannabis* showed short and thin plant type and short life cycle when they were transplanted to lower latitude (Zhang et al., 2018). Similarly, *Z. latifolia* individuals from the north regions, especially from HLR and LHR regions, exhibited thin stem and short plant height. Furthermore, the north *Z. latifolia* populations initially entered flowering in the common garden because the shortened day length could promote pre-flowering. Conversely, the prolonged daylight hours could delay the onset of sexual reproduction in plants. Indeed, the individuals from the south-most region ZJR did not bloom in the common garden. This suggested that, although low latitude originated plant populations are often expected to migrate to higher latitude under warming temperature(Walther et al., 2002; Deutsch et al., 2008), *Z. latifolia* may not have sexual reproduction when they migrate to higher latitude, implying the meaningless of migration which cannot effectively occupy specific habitats.

### The Relationships among Genetic, Geographic and Growth Traits

Both genotype and environment determine intraspecific variations which allow the species to deal with the ever-changing climates (Nicotra et al., 2010). In this study, IBD (isolation by distance) model within current *Z. latifolia* populations was manifested by the significant positive correlation between genetic distance and geographic distance, which was agreeable to previous studies of *Z. latifolia*(Chen et al., 2012; Chen et al., 2017). Additionally, IBE (isolation by environment) model for *Z. latifolia* populations was suggested by the following evidences: (i) The morphological traits of populations in common garden were positively correlated with temperatures of their original habitats; (ii) The degree of differentiation of morphological traits among populations (*Q*_ST_=0.931-0.998) were much greater than the genetic differentiation estimated by SSRs (*F*_ST_= 0.585); (iii) In the common garden located in the Yangtze River Basin, the populations from the Yangtze River region showed better fitness than populations from other regions. Therefore, both IBD and IBE models influenced the genetic patterns of *Z. latifolia*, like another emergent macrophyte *Phragmites australis* (Ren et al., 2020).

Furthermore, the phenotypic plasticity in *Z. latifolia* probably has a genetic basis considering the significant correlation between growth traits and genetic distance. The heritable phenotypic plasticity could thus imply the potential crisis of decreasing in adaptation or competitiveness when environment changed swiftly since it was encoded in the genome and couldn’t change as rapidly as environments did.

## CONCLUSIONS

Global warming is the rapid increase surface average temperature of Earth’s over the past century. The effects of the global warming on biosphere still need to be explored, including the effects on the plant species. In this study, we chose the East Asia widespread aquatic plant *Z. latifolia* to investigate the global warming on the growth traits, reproduction traits and the photosynthesis traits in a common garden. We found that the higher latitude populations grew compact with shorter and narrower leaf, thinner stem and less biomass. All the changes of growth traits indicated that migrators from high to low latitudes could be extinct when they competed to other species under warmer temperature. The lower latitude originated populations didn’t perform sexual reproduction when grew at higher latitude. The result suggests that, under warmer temperature, the lower latitude *Z. latifolia* populations don’t have the capacity to occupy higher latitude habitats in the long term. Therefore, all our results suggested that *Z. latifolia* populations could not adapt to the global warming and would be at least at risk of local extinction in a warmer future.

## DATA AVAILABILITY STATEMENT

The datasets presented in this study can be found in online repositories. The names of the repository/repositories and accession number(s) can be found in the article Supplementary Material.

## AUTHOR CONTRIBUTIONS

HJ, XF, YC, and WL designed the research. XF and YC collected populations in field. XF, YC, GKW and HJ performed experiments. HJ, XF, YC analyzed the data and wrote the manuscript. All authors read and approved the final manuscript.

## ACKNOWLEDGEMENT/FUNDING

This work was supported by the Strategic Priority Research Program of Chinese Academy of Sciences (grant number XDB31000000) and by the Youth Innovation Promotion Association CAS (2021340). We would like to thank Dr. Wenlong Fu for plant collecting in the field and to thank Mr. Zhenwei Lu for the *Z. latifolia* transplanting in the common garden.

## Supporting information

**Figure S1.**
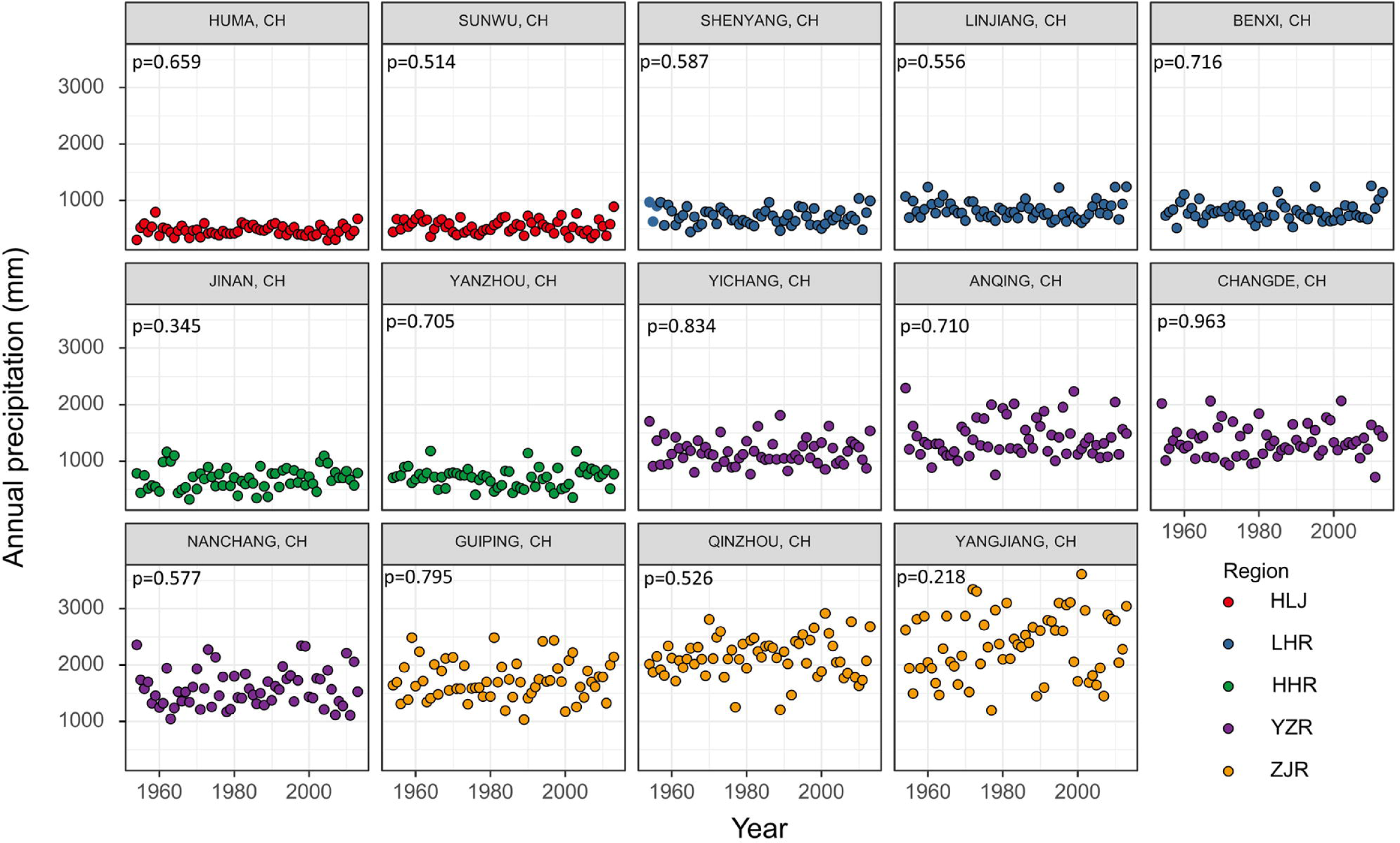
Changes of annual precipitation at sampling sites in the past 60 years (1954-2014).

Table S1. Sampling information of *Zizania latifolia*.

Table S2. SSR primer sequences used in this study.

Table S3. Information of meteorological stations

Table S4. Relationship between growth and reproduction traits of *Z. latifolia* in common garden and their original habitat yearly average temperature and annual precipitation

Table S5. Estimated trait differentiation *Q*_ST_ in *Z. latifolia*.

