## Supplemental Tables for "Latitude-Driven Functional Trait Variations in *Zizania latifolia*: Insights into Climate Adaptation"

Table S1. Sampling information of *Zizania latifolia*.

| Region | Population code | Location | Latitude (N) | Longitude (E) | Individual number | |
| --- | --- | --- | --- | --- | --- | --- |
|  |  |  |  |  | Total*^a^* | 2017*^b^* |
| Heilongjiang River Basin (HLJ) | BSLZ | Baishilizi | 50°54′14.76″ | 127°8′1.30″ | 22 | 14 |
|  | XXT | Xiaoxintun | 50°44′18.49″ | 127°15′47.73″ | 21 | 10 |
|  | LQQ | Lanqiqiao | 49°54′20.00″ | 127°29′38.2″ | 20 | 14 |
|  | HDY | Hadayan | 49°33′39.6″ | 127°55′33.8″ | 20 | 14 |
|  | HW | Hongwei | 49°29′33.2″ | 128°40′2.2″ | 20 | 14 |
|  | KEB | Kuerbin | 49°23′18.3″ | 128°56′56.4″ | 22 | 14 |
|  | YLZ | Yuliangzi | 49°12′12.8″ | 129°43′28.3″ | 21 | 14 |
| Liaohe River Basin (LHR) | LZ | Liaozhong | 42°23′15.5″ | 125°46′4.9″ | 20 | 14 |
|  | JH | Jinhua | 41°56′6.4″ | 122°52′38.6″ | 20 | 14 |
|  | HR | Huanren | 41°55′28.22″ | 124°04′58.18″ | 20 | 14 |
|  | DG | Donggang | 41°32′26.5″ | 122°44′23.3″ | 20 | 14 |
|  | ZD | Zhangdang | 41°14′35.4″ | 125°21′51.5″ | 20 | 14 |
| Huanghe River Basin (HHR) | HXD | Huanxiang-dian | 36°45′6.18″ | 117°6′29″ | 20 | 14 |
|  | MK | Mankou | 35°58′50.05″ | 116°15′28.05″ | 20 | 14 |
|  | DP | Dongping | 35°11′25.5″ | 116°40′56.28″ | 20 | 14 |
|  | LQ | Luqiao | 34°58′9.4″ | 116°54′52.7″ | 20 | 14 |
| Yangtze River Basin (YZR) | LZL | Liangzi lake | 30°50′39.4″ | 117°20′53.1″ | 21 | 14 |
|  | LGL | Longgan lake | 30°24′26.7″ | 112°28′44.4″ | 21 | 14 |
|  | SJL | Shengjin lake | 30°24′7.5″ | 117°02′51.5″ | 20 | 14 |
|  | BDL | Baidang lake | 30°06′11.9″ | 114°27′29.3″ | 22 | 14 |
|  | CHL | Changhu lake | 30°03′46″ | 115°56′54.6″ | 21 | 14 |
|  | DTL | Dongting lake | 29°46′27.5″ | 113°14′36″ | 21 | 14 |
|  | HHL | Honghu lake | 29°30′15.5″ | 112°47′10.1″ | 21 | 14 |
| Zhujiang River Basin (ZJR) | DC | Dongcheng | 22°38′17.7″ | 108°22′50.7″ | 21 | 14 |
|  | WC | Wuchuan | 22°42′14.1″ | 110°21′41.9″ | 21 | 14 |
|  | BL | Beiliu | 22°22′47.2″ | 112°37′46.2″ | 23 | 14 |
|  | FC | Fangcheng-gang | 21°40′3.9″ | 108°17′45.5″ | 21 | 14 |
|  | NM | Nama | 21°20′58.9″ | 110°38′10.0″ | 21 | 14 |

*a*: number of total individuals collected from field

*b*: number of individuals sampled for growth and reproduction traits measurement in 2017

Table S2. SSR primer sequences used in this study.

| Primer Name | Sequence (5’ – 3’) | Repeat motif | species | Reference |
| --- | --- | --- | --- | --- |
| ZM4 | F: AGCAGGAAGCAACTGACTTA | AG | *Z. latifolia* | Quan et al., 2009 |
|  | R: GCATCTCAGGGTGAAAGAAC |  |  |  |
| ZM25 | F: GTTCTGAGTTGCAACCTGGT | CA, TA | *Z. latifolia* | Quan et al., 2009 |
|  | R: CCCATATGTCAGCGAGACAT |  |  |  |
| ZM26 | F: CGAACCCTGCATCAAACACT | AG | *Z. latifolia* | Quan et al., 2009 |
|  | R: GATTCGGGAGTCTCCTAGTT |  |  |  |
| ZM28 | F: CCCTTGCTCATGCATAGATG | GA, AG | *Z. latifolia* | Quan et al., 2009 |
|  | R: CACCTTGCACATCAGCTCAT |  |  |  |
| ZM35 | F: GACTGATGACAACTGATGGA | GA | *Z. latifolia* | Quan et al., 2009 |
|  | R: GCACATGCTTGTGTACTTGT |  |  |  |
| ZM44 | F: TGCGTGATCTTCAATTCCAA | GA | *Z. latifolia* | Quan et al., 2009 |
|  | R: GCTACCGAAGATGTCGTTG |  |  |  |
| RM14233 | F: CAGCAAATAACCCTTCGTCATGG | AC | *O. sativa* | Wang et al., 2015 |
|  | R: CAGCACTGCTCTGGATTCTTTGG |  |  |  |
| RM20188 | F: AGGCGACAGGGACACGATGG | CCG | *O. sativa* | Wang et al., 2015 |
|  | R: AAGCCGGATATTCCGTGTGAAGG |  |  |  |
| RM28090 | F: ATCGATCCTCAAGGCAGCATGG | AGC | *O. sativa* | Wang et al., 2015 |
|  | R: GGCTTGGAAGTTCAGGCACAGG |  |  |  |
| Zt1 | F: GCAAATCTCCTGTCTTTTTCT | TAGA | *Z. texana* | Richards et al., 2007 |
|  | R: GTTTAGCCAGCTCCCAATGTA |  |  |  |
| Zt23 | F: GGACGTTGACATTTTCACA | AG | *Z. texana* | Richards et al., 2007 |
|  | R: GGATCAGTAAATCCAAATCTGT |  |  |  |
| ZL1 | F: GCTGCTACAATCTTTCTAACTACTA | TA | *Z. latifolia* | Wagutu et al., 2020 |
|  | R: TTCCAGGCTGCGTTTT |  |  |  |
| ZL3 | F: TTTGATGCTGCTACAATCTTTC | TA | *Z. latifolia* | Wagutu et al., 2020 |
|  | R: TTCCAGGCTGCGTTTT |  |  |  |
| ZL4 | F: GCTGCTACAATCTTTCTAACTACTA | TA | *Z. latifolia* | Wagutu et al., 2020 |
|  | R: TTCCAGGCTGCGTTTT |  |  |  |
| ZL5 | F: GTTCTTTGATGCTGCTACAAT | TA | *Z. latifolia* | Wagutu et al., 2020 |
|  | R: TTCCAGGCTGCGTTTT |  |  |  |
| ZL9 | F: CATTGCCACTAGACATACA | AT | *Z. latifolia* | Wagutu et al., 2020 |
|  | R: TTGAGGATTCGACGATA |  |  |  |
| ZL10 | F: ATCATTGCCACTAGACATAC | AT | *Z. latifolia* | Wagutu et al., 2020 |
|  | R: CTTGATGACAAAGGATAGAA |  |  |  |
| ZL31 | F: TTGAGGATCAGGTGGCAGTC | CT | *Z. latifolia* | Wagutu et al., 2020 |
|  | R: TGACCATTCAGCTTCTTGGA |  |  |  |
| ZL32 | F: TTGAGGATCAGGTGGCAGTC | CT | *Z. latifolia* | Wagutu et al., 2020 |
|  | R: ACCATTCAGCTTCTTGGAGA |  |  |  |
| ZL36 | F: CATGCTTCTCATCGGTAGAG | AT | *Z. latifolia* | Wagutu et al., 2020 |
|  | R: GTGACACCAAACAATGTCAA |  |  |  |
| ZL42 | F: TTGTTACGAGGACTTTATGAG | AT | *Z. latifolia* | Wagutu et al., 2020 |
|  | R: ATTCTGTACGTTTGAGGTTGT |  |  |  |
| ZL43 | F: TTGTTACGAGGACTTTATGAG | AT | *Z. latifolia* | Wagutu et al., 2020 |
|  | R: TGATTCTGTACGTTTGAGGTT |  |  |  |
| Zl55 | F: GTTTGAGACGGCTGTTTTG | TC | *Z. latifolia* | Wagutu et al., 2020 |
|  | R: CAGGAGGCATGAGGAAGG |  |  |  |
| ZL56 | F: GTTTGAGACGGCTGTTTTG | TC | *Z. latifolia* | Wagutu et al., 2020 |
|  | R: AGCAGGAGGCATGAGGAA |  |  |  |
| ZL57 | F: GTTTGAGACGGCTGTTTTG | TC | *Z. latifolia* | Wagutu et al., 2020 |
|  | R: AGAGCAGGAGGCATGAGG |  |  |  |

**Quan Z, Pan L, Ke W, Liu Y, Ding Y. 2009.** Sixteen polymorphic microsatellite markers from *Zizania latifolia* Turcz. (Poaceae). *Molecular Ecology Resources* **9**: 887–889.

**Richards CM, Antolin MF, Reilley A, Poole J, Walters C. 2007.** Capturing genetic diversity of wild populations for ex situ conservation: Texas wild rice (*Zizania texana*) as a model. *Genetic Resources and Crop Evolution* **54**: 837 – 848.

**Wagutu GK, Njeri HK, Fan XR, Chen YY. 2020.** Development and transferability of SSR primers in the wild rice *Zizania latifolia* (Poaceae). *Plant Science Journal* **38**: 105–111.

**Wang HM, Wu GL, Jiang SL, Huang QN, Feng BH, Hunag CG, Jiang SM, Wu JL. 2015.** Genetic diversity of *Zizania latifolia* Griseb. from Poyang lake basin based on SSR and ISSR analysis. *Journal of Plant Genetic Resources* **16**: 133–141.

Table S3. Information of meteorological stations

| Region | Meteorological station | Latitude | Longitude |
| --- | --- | --- | --- |
| HLJ | HUMA, CH | 50°43.02′ | 126°39′ |
|  | SUNWU, CH | 49°25.98′ | 127°21′ |
| LHR | SHENYANG, CH | 41°43.98′ | 123°31.02′ |
|  | LINJIANG, CH | 41°43.02′ | 126°55.02 |
|  | BENXI, CH | 41°19.02′ | 123°46.98′ |
| HHR | JINAN, CH | 36°36′ | 117°3′ |
|  | YANZHOU, CH | 35°34.02′ | 116°51′ |
| YZR | YICHANG, CH | 30°42′ | 111°18′ |
|  | ANQING, CH | 30°31.98′ | 117°3′ |
|  | CHANGDE, CH | 29°3′ | 111°40.98′ |
|  | NANCHANG, CH | 28°36′ | 115°55.02′ |
| ZJR | GUIPING, CH | 23°24′ | 110°4.98′ |
|  | QINZHOU, CH | 21°57′ | 108°37.02′ |
|  | YANGJIANG, CH | 21°52.02′ | 111°58.02′ |

Table S4. Relationship between growth and reproduction traits of *Z. latifolia* in common garden and their original habitat yearly average temperature and annual precipitation

| Traits | Model^a^ | R^2^ | Regression coefficient | *p* value |
| --- | --- | --- | --- | --- |
| Tiller number | Y= a_0_+a_1_X_1_+a_2_X_1_^2^+ a_3_X_2_ | 0.356 | a_0_=11.76 | <10^-13^ |
|  |  |  | a_1_=2.135 | <10^-16^ |
|  |  |  | a_2_=-0.186 | <10^-16^ |
|  |  |  | a_3_=0.022 | <10^-16^ |
| Plant height | Y=a_0_+a_1_X_1_+a_2_X_2_+a_3_X_1_:X_2_ | 0.855 | a_0_=46.25 | <10^-16^ |
|  |  |  | a_1_=4.65 | <10^-16^ |
|  |  |  | a_2_=0.024 | 0.0026 |
|  |  |  | a_3_=-0.0017 | 6×10^-7^ |

| Leaf length | Y=a_0_+a_1_X_1_+a_2_X_2_+a_3_X_1_:X_2_ | 0.8841 | a_0_=26.22 | <10^-16^ |
| --- | --- | --- | --- | --- |
|  |  |  | a_1_=2.81 | <10^-16^ |
|  |  |  | a_2_=0.005 | 0.27 |
|  |  |  | a_3_=-0.0005 | 0.013 |

| Leaf width | Y=a_0_+a_1_X_1_+a_2_X_2_+a_3_X_1_:X_2_ | 0.8854 | a_0_=1.19 | <10^-16^ |
| --- | --- | --- | --- | --- |
|  |  |  | a_1_=0.39 | <10^-16^ |
|  |  |  | a_2_=-0.007 | <10^-8^ |
|  |  |  | a_3_=0.0003 | <10^-12^ |
| Internode number | Y=a_0_+a_1_X_1_+a_2_X_2_+a_3_X_1_:X_2_ | 0.179 | a_0_=2.73 | 7×10^-12^ |
|  |  |  | a_1_=0.079 | 5×10^-8^ |
|  |  |  | a_2_=0.0025 | 3×10^-4^ |
|  |  |  | a_3_=-0.00014 | 2×10^-7^ |
| Internode length | Y= a_0_+a_1_X_1_+a_2_X_1_^2^+ a_3_X_2_ | 0.143 | a_0_=3.80 | <10^-16^ |
|  |  |  | a_1_=0.39 | <10^-12^ |
|  |  |  | a_2_=-0.024 | <10^-9^ |
|  |  |  | a_3_=-0.0014 | 0.047 |
| Dry weight | Y=a_0_+a_1_X_1_+a_2_X_2_ | 0.7631 | a_0_=5.73 | 0.0001 |
|  |  |  | a_1_=2.42 | <10^-16^ |
|  |  |  | a_2_=0.008 | 0.0003 |
| Diameter | Y=a_0_+a_1_X_1_+a_2_X_2_+a_3_X_1_:X_2_ | 0.9525 | a_0_=8.23 | <10^-16^ |
|  |  |  | a_1_=0.37 | <10^-16^ |
|  |  |  | a_2_=-0.009 | <10^-16^ |
|  |  |  | a_3_=0.0004 | <10^-16^ |
| Spike frequency | Y= a_0_+a_1_X_1_^2^ | 0.5785 | a_0_=85.57 | <10^-11^ |
|  |  |  | a_1_=-0.186 | <10^-6^ |
| Spike number | Y= a_0_+a_1_X_1_+a_2_X_1_^2^+ a_3_X_2_ | 0.5271 | a_0_=-0.114 | 0.677 |
|  |  |  | a_1_=0.353 | <10^-16^ |
|  |  |  | a_2_=-0.039 | <10^-16^ |
|  |  |  | a_3_=0.0049 | <10^-16^ |
| Spike length | Y=a_0_+a_1_X_1_+a_2_X_2_ | 0.7658 | a_0_=14.96 | <10^-16^ |
|  |  |  | a_1_=1.61 | <10^-16^ |
|  |  |  | a_2_=-0.0007 | 0.77 |
| Grain per spike | Y=a_0_+a_1_X_1_+a_2_X_2_+a_3_X_1_:X_2_ | 0.6346 | a_0_=301 | <10^-6^ |
|  |  |  | a_1_=9.25 | 0.008 |
|  |  |  | a_2_=-0.46 | <10^-5^ |
|  |  |  | a_3_=0.024 | <10^-4^ |

^a^: Y: growth traits or reproduction trats; X_1_: temperature (℃); X_2_: precipitation (mm)

Table S5. Estimated trait differentiation *Q*_ST_ in *Z. latifolia*.

| Trait | V_among_ | V_within_ | *Q*_ST_ |
| --- | --- | --- | --- |
| Height | 11525 | 63 | 0.995 |
| Internode number | 9.505 | 0.461 | 0.954 |
| Internode length | 73.9 | 1.63 | 0.978 |
| Tiller number | 1108 | 35.3 | 0.969 |
| Diameter | 305.7 | 0.54 | 0.998 |
| Leaf length | 5345 | 34 | 0.994 |
| Leaf width | 348.2 | 2.3 | 0.993 |
| Dry weight | 9351 | 141 | 0.985 |
| Spike number | 55.6 | 0.91 | 0.984 |
| Spike frequency*^a^* | 9891 | 484 | 0.953 |
| Grain per spike | 132398 | 9830 | 0.931 |
| Spike length | 2018 | 75.9 | 0.964 |

*a*: the *Q*_ST_ of spike frequency estimated by variance among and within regions to instead of populations because there was only one spike frequency for each population that was impossible to estimate the variance within population.
